# When does high-dose antimicrobial chemotherapy prevent the evolution of resistance?

**DOI:** 10.1101/020321

**Authors:** Troy Day, Andrew F. Read

## Abstract

High-dose chemotherapy has long been advocated as a means of controlling drug resistance in infectious diseases but recent empirical and theoretical studies have begun to challenge this view. We show how high-dose chemotherapy engenders opposing evolutionary processes involving the mutational input of resistant strains and their release from ecological competition. Whether such therapy provides the best approach for controlling resistance therefore depends on the relative strengths of these processes. These opposing processes lead to a unimodal relationship between drug pressure and resistance emergence. As a result, the optimal drug dose always lies at either end of the therapeutic window of clinically acceptable concentrations. We illustrate our findings with a simple model that shows how a seemingly minor change in parameter values can alter the outcome from one where high-dose chemotherapy is optimal to one where using the smallest clinically effective dose is best. A review of the available empirical evidence provides broad support for these general conclusions. Our analysis opens up treatment options not currently considered as resistance management strategies, and greatly simplifies the experiments required to determine the drug doses which best retard resistance emergence in patients.

**Significance Statement:** The evolution of antimicrobial resistant pathogens threatens much of modern medicine. For over one hundred years, the advice has been to ‘hit hard’, in the belief that high doses of antimicrobials best contain resistance evolution. We argue that nothing in evolutionary theory supports this as a good rule of thumb in the situations that challenge medicine. We show instead that the only generality is to either use the highest tolerable drug dose or the lowest clinically effective dose; that is, one of the two edges of the therapeutic window. This approach suggests treatment options not currently considered, and greatly simplifies the experiments required to identify the dose that best retards resistance evolution.

---

Antimicrobial resistance is one of greatest challenges faced by modern medicine. There is a widely held view that the evolutionary emergence of drug resistance is best slowed by using high doses of drugs to eliminate pathogens as early and quickly as possible. This view, first expounded by Ehrlich (1) (‘hit hard’) and later Fleming (2) (‘if you use penicillin, use enough’), is today encapsulated in the advice to administer ‘the highest tolerated antibiotic dose’ (3, 4). The rationale is two-fold. First, a high concentration of drug will eliminate drug-sensitive microbes quickly and thereby limit the appearance of resistant strains. Second, a high concentration of drug will also eliminate strains that have some partial resistance, provided the concentration is above the so-called mutant prevention concentration (MPC) (5–12).

This is an intuitively appealing idea, but several authors have recently questioned whether high-dose chemotherapy is, as a generality, defensible in terms of evolutionary theory (13–16). This is because the use of extreme chemical force comes at the cost of maximizing the selective advantage of the very pathogens that we fear most; namely, those which cannot be eradicated by safely administered doses of drug. Some experimental studies have also shown that lighter-touch chemotherapy not only better prevents the emergence of resistance but it restores host health just as well as high-dose chemotherapy (15–17).

Here we examine when high-dose chemotherapy is the best strategy and when it is not, by developing a general mathematical model for resistance emergence within a treated patient using principles from evolutionary biology. The analysis shows that high-dose chemotherapy gives rise to opposing evolutionary processes. As a result, the optimal therapy for controlling resistance depends on the relative strengths of these processes. High-dose therapy can, in some circumstances, retard resistance emergence but evolutionary theory provides no support for using this strategy as a general rule of thumb, nor does it provide support for focussing on the MPC as a general approach for resistance prevention. More broadly we find that the opposing evolutionary processes lead to a unimodal relationship between drug concentration and resistance emergence. Therefore the optimal strategy is to use either the largest tolerable dose or the smallest clinically effective dose. We illustrate these general points with a simple model that shows how a seemingly minor change in parameter values can alter the outcome from one where high-dose chemotherapy is optimal to one where using the smallest clinically effective dose is best. A review of the empirical evidence provides broad support for these conclusions.

## A Theoretical Framework for Resistance Evolution

Determining a patient treatment regimen involves choosing an antimicrobial drug (or drugs) and determining the frequency, timing, and duration of administration. The impact of each of these on resistance emergence has been discussed elsewhere (e.g., 9, 18). Here we focus solely on drug concentration because it has historically been the factor most often discussed, and because it is the source of recent controversy (e.g., 10, 12–14, 16). We seek to understand how the probability of resistance emergence changes as a function of drug concentration.

For simplicity we assume that drug concentration is maintained at a constant level during treatment and refer to this concentration as ‘dose’. This assumption is not meant to be realistic but it serves as a useful tool for gaining a better understanding of how drug resistance evolves. After laying the groundwork for this simple case we show in the Appendix that allowing for more realistic pharmacokinetics does not alter our qualitative conclusions.

Drug resistance is a matter of degree, with different genotypes having different levels of resistance (measured, for example, as the minimum inhibitory concentration, MIC). Our main focus is on what we call high-level resistance (HLR). This will be defined precisely below but for the moment it can be thought of as resistance that is high enough to render the drug ineffective (so that its use is abandoned). We begin by supposing that the HLR strain is one mutational step away from the wild type but we relax this assumption in the Appendix.

Why is it that resistant strains reach appreciable densities in infected patients only once drug treatment is employed? The prevailing view is that there is a cost of resistance in the absence of the drug, but that this cost is compensated for by resistance in the presence of the drug. It is not the presence of the drug *per se* that provides this compensation; rather, it is the removal of the wild type by the drug that does so (13, 19). This implies that the presence of the wild type competitively suppresses the resistant strain, and that drugs result in the spread of such strains because they remove this competitive suppression (a process called ‘competitive release’; 19).

To formalize these ideas, consider an infection in the absence of treatment. The wild type pathogen enters the host and begins to replicate. As it does so, it consumes resources and stimulates an immune response. We use *P*(*t*) to denote the density of the wild type and *X*(*t*) to denote a vector of within-host state variables (e.g., density of immune system components, resources, etc). Without loss of generality we suppose that the vector *X* is defined in such a way that pathogen replication causes its components to decrease. This decrease in *X*, in turn, makes the within-host environment less favorable for pathogen replication. If *X* is suppressed enough, the net replication rate of the wild type will reach zero. Thus *X* can be viewed as the quality of the within-host environment from the standpoint of pathogen replication.

As the wild type replicates it gives rise to the HLR strain through mutation and the initial infection might include some HLR pathogens as well. But the HLR strain is assumed to bear some metabolic or replicative cost, meaning that it is unable to increase in density once the wild type has become established. Mechanistically this is because the wild type suppresses the host state, *X*, below the minimum value required for a net positive replication by the HLR strain (19). Thus, we ignore the effect of the HLR strain when modeling the joint dynamics of *P*(*t*) and *X*(*t*) in the absence of treatment (see Appendix A for details).

At some point (e.g., the onset of symptoms) drug treatment is introduced. Provided the dosage is high enough the wild type will be driven to extinction. We use *c* to denote the (constant) concentration of the drug in the patient. We distinguish between *theoretically possible* versus *feasible* doses. Theoretically possible doses are those that can be applied *in vitro*. Feasible doses are those that can, in practice, be used *in vivo*. There will be a smallest clinically effective dose that places a lower bound on the feasible values of *c* (denoted *c*_*L*_) and a maximum tolerable dose because of toxicity (denoted *c*_*U*_). The dose range between these bounds is called the therapeutic window (20).

Once treatment has begun, we use *p*(*t*; *c*) and *x*(*t*; *c*) to denote the density of the wild type strain and the within-host state. This notation reflects the fact that different dosages (i.e., concentrations) will give rise to different trajectories of *p* and *x* during the remainder of the infection. We model the dynamics of *p* and *x* deterministically during this phase.

As the wild type is driven to extinction it will continue to give rise to HLR microbes through mutation. The mutation rate is given by a function *λ*[*p*(*t*; *c*), *c*] that is increasing in *p* and decreasing in *c*. We suppose that lim_*c→∞*_ *λ*[*p, c*] = 0 because a high enough drug concentration will completely suppress wild type replication and thus mutation. Any HLR microbes that are present during treatment will no longer be destined to rarity because they will be released from competitive suppression (19). We use *π*[*x*(*t*; *c*), *c*] to denote the probability of escaping initial extinction when rare. The function *π* is increasing in *x* because it is through this state that the HLR strain has been competitively suppressed (19). And *π* is decreasing in *c* with lim_*c→∞*_ *π*[*x, c*] = 0 because a high enough dose will also suppress even then HLR strain.

We can now provide a precise definition of high-level resistance (HLR). Although lim_*c→∞*_ *π*[*x, c*] = 0, the concentration at which this limit is reached can lie outside the therapeutic window [*c*_*L*_, *c*_*U*_]. We define HLR to mean that *π*[*x, c*] is very nearly equal to *π*[*x,* 0] over the therapeutic window. Biologically this means that, in terms of clinically acceptable doses, significant suppression of HLR is not possible. We focus on HLR because, for genotypes that do not satisfy this property, there is then no resistance problem to begin (since one can always use a high enough dose to remove all pathogens).

With the above formalism, we focus on resistance emergence, defined as the replication of resistant microbes to a high enough density within a patient to cause symptoms and/or to be transmitted (19). In the analytical part of our results this is equivalent to the resistant strain not being lost by chance while rare.

The probability of resistance emergence is approximately equal to 1 − *e*^−*H*(*c*)^ where

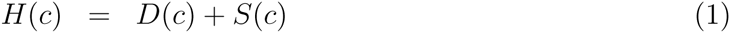

and

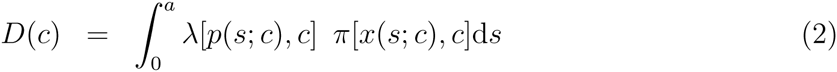

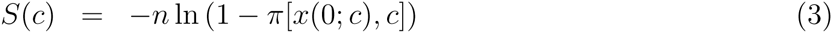

(see Appendix A). We refer to *H*(*c*) as the resistance ‘hazard’, and *a* is the duration of treatment with *s* = 0 corresponding to the start of treatment. The quantity *D*(*c*) is the *de novo* hazard - it is the hazard due to resistant strains that appear *de novo* during treatment. It is comprised of the integral of the product of *λ*[*p*(*s*; *c*), *c*], the rate at which resistant mutants appear at time *s* after the start of treatment, and *π*[*x*(*s*; *c*), *c*], the probability of escape of any such mutant. The quantity *S*(*c*) is the standing hazard - it is the hazard due to a standing population of *n* resistant microbes that are already present at the beginning of treatment (see Appendix A). To minimize the probability of resistance emergence we therefore want to minimize the hazard *H*(*c*), subject to the constraint that the dosage *c* falls within the therapeutic window [*c*_*L*_, *c*_*U*_].

## Results

To determine how high-dose chemotherapy affects the probability of resistance emergence we determine how *H*(*c*) changes as drug dosage *c* increases. Differentiating expression (1) with respect to *c* we obtain

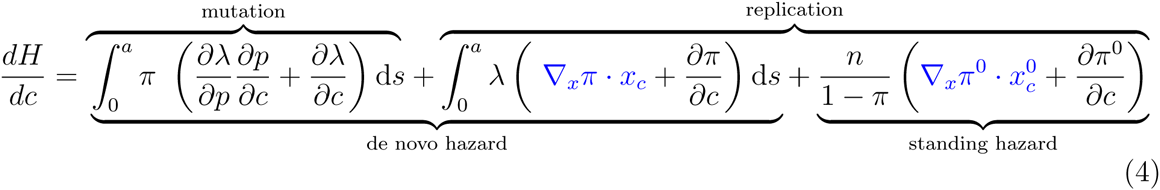

where *π*^0^ = *π*[*x*(0; *c*), *c*], *x*^0^ = *x*(0; *c*), and subscripts denote differentiation. Equation (4) is partitioned in two different ways to better illustrate the effect of increasing dose. The first is a partitioning of its effect on mutation and replication. The second is a partitioning of its effect on the de novo and standing hazards. We have also indicated the terms that represent competitive release in blue (as explained below).

The first term in equation (4) represents the change in *de novo* mutation towards the HLR strain that results from an increase in dose. The term (*∂λ/∂p*)(*∂p/∂c*) is the change in mutation rate, mediated through a change in wild type density; *∂λ/∂p* specifies how mutation rate changes with an increase in the wild type density *p* (positive) while *∂p/∂c* specifies how the wild type density changes with an increase in dose (typically negative for much of the duration of treatment). Thus the product, when integrated over the duration of treatment, is expected to be negative. The term *∂λ/∂c* is the change in mutation rate that occurs directly as a result of an increased dose (e.g., the direct suppression of wild type replication, which suppresses mutation). This, is expected to be non-positive in the simplest cases and is usually taken as such by proponents of high-dose chemotherapy. Therefore *high-dose chemotherapy decreases the rate at which HLR mutations arise during treatment.* Note, however, that if the drug itself causes a higher mutation rate (e.g., 21), then it is possible for an increased dose to increase the rate at which resistance appears.

The second term in equation (4) represents replication of HLR strains once they have appeared *de novo* during the course of treatment. The term ∇_*x*_*π · x*_*c*_ is the indirect increase in escape probability, mediated through the effect of within-host state, *x*. Specifically, *x*_*c*_ is a vector whose elements give the change in each state variable arising from an increased dosage (through the removal of the wild type). These elements are typically expected to be positive for much of the duration of treatment because an increase in dose causes an increased rebound of the within-host state through a heightened removal of wild type microbes. The quantity ∇_*x*_*π* is the gradient of the escape probability with respect to host state *x*, and its components are expected to be positive (higher state leads to a greater probability of escape). The integral of the dot product ∇_*x*_*π · x*_*c*_ is therefore the competitive release of the HLR strain in terms of de novo hazard (19). This will typically be positive. The term *∂π/∂c* is the direct change in escape probability of de novo mutants as a result of an increase in dosage (i.e., the extent to which the drug suppresses even the HLR strain). This term is negative at all times during treatment but, by the definition of HLR, this is small. Therefore, *high-dose chemotherapy increases the replication of any HLR mutants that arise de novo during treatment.*

Finally, the third term in equation (4) represents the replication of any HLR strains that are already present at the start of treatment. The term 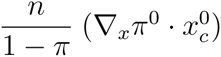 is the indirect effect of dose on standing hazard, where *n* is the number of resistant pathogens present at the start of treatment. The quantity 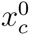 is again a vector whose elements give the change in state arising from increased dosage (through the removal of the wild type). The components of this are typically expected to be positive because an increase in dose causes a rebound in the within-host state. ∇_*x*_*π*^0^ is the gradient of the escape probability with respect to state, and its components are expected to be positive (higher state leads to greater probability of escape). The dot product of the two, ∇_*x*_*π · x*_*c*_, is therefore the competitive release of the HLR strain in terms of standing hazard (19). This will typically be positive. The term 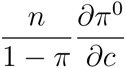 is the direct change in escape probability of pre-existing mutants as a result of an increase in dosage (i.e., the extent to which the drug suppresses even these HLR mutants) and is negative. Again, however, by the definition of HLR, this will be small and therefore *high-dose chemotherapy increases the replication of any HLR mutants that are present at the start of treatment.* Appendix B shows that the same set of qualitative factors arise if there are strains with intermediate resistance as well.

The above results provide a mathematical formalization of earlier verbal arguments questioning the general wisdom of using high-dose chemotherapy as a means of controlling resistance emergence (13, 16). Advocates of the conventional heavy dose strategy tend to emphasize how high-dose chemotherapy can reduce mutational input and potentially even suppress the replication of resistant strains (the black derivatives in equation 4). However, high-dose chemotherapy leads to competitive release and thus greater replication of any resistant strains that are present (the blue derivatives in equation 4). Equation (4) shows that it is the relative balance among these opposing processes that determines whether high-dose chemotherapy is the optimal approach. We will present a specific numerical example shortly that illustrates these points, but first we draw two more general conclusions from the theory.

### (1) Intermediate doses yield the largest hazard and thus the greatest likelihood of resistance emergence across all theoretically feasible doses

The opposing evolutionary processes explained above are the reason for this result (also see 16). First note that the functions *λ* and *π* will typically be such that *D*(0) *≈* 0. In other words, the HLR strain does not emerge *de novo* within infected individuals if they are not receiving treatment. Mechanistically, this is because any resistant strains that appear tend to be competitively suppressed by the wild type strain (19). Although, *S*(0) need not be zero (see Appendix C, Figure C2), the rate of change of *S*(*c*) with respect to *c* (i.e., the third term in equation 4) is positive at *c* = 0. Therefore the maximum hazard cannot occur at *c* = 0.

Second, for large enough doses we have *π*[*x*(*s*; *c*), *c*] *≈* 0 for all *s* because such extreme concentrations will prevent replication of even the HLR strain. This makes both the *de novo* hazard *D*(*c*) and the standing hazard *S*(*c*) zero. Furthermore, for large enough *c* we also have *λ*[*p*(*s*; *c*), *c*] *≈* 0 for all *s* as well if HLR can arise only during wild type replication, because such extreme concentrations prevent all replication of the wild type. This is an additional factor making the *de novo* hazard *D*(*c*) decline to zero for large *c*. Therefore lim_*c→∞*_ *H*(*c*) = 0 and so the maximum hazard cannot occur for large values of *c* either (16). Thus, the maximum hazard must occur for an intermediate drug dosage. Although this prediction is superficially similar to that of the mutant selection window hypothesis (5–9), there are important differences between the two as will be elaborated upon in the discussion.

### (2) The optimal dose is either the maximum tolerable dose or minimum clinically effective dose

We have seen that the maximum hazard occurs for an intermediate dose. Suppose, further, that the hazard *H*(*c*) is a unimodal function of *c* (i.e., it has a single maximum). Several specific mathematical models (Day unpubl. results) and a body of empirical work (see Discussion) are consistent with that assumption. Then the drug dose which best reduces the probability of resistance emergence is always at one of the two extremes of the therapeutic window. This means that it is best to use either the smallest clinically effective dose or the largest tolerable dose depending on the situation, but never anything in between (Figure 1).

**Figure 1.**
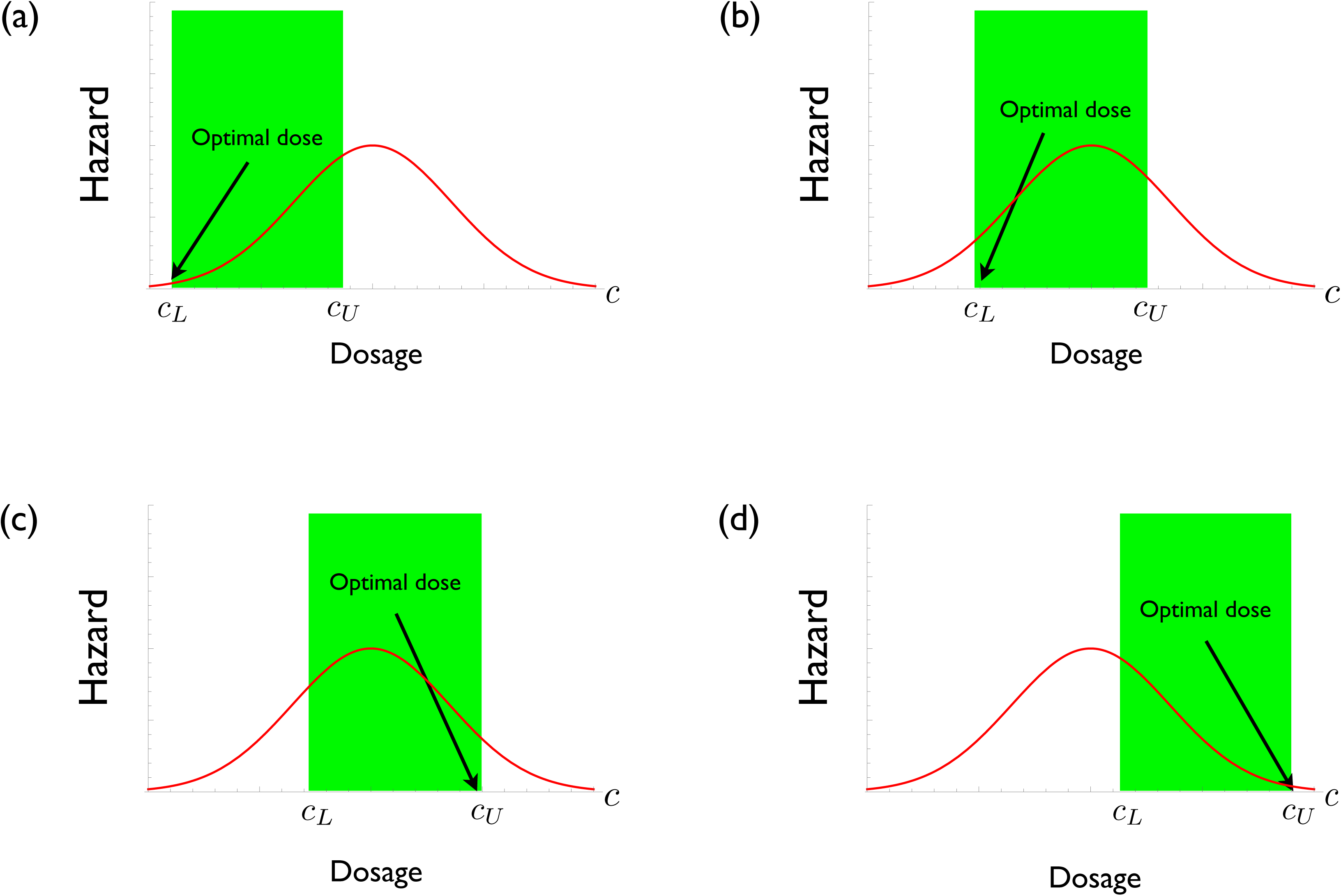
Hypothetical plots of resistance hazard *H*(*c*) as a function of drug concentration *c*, with the lowest effect dose and the highest tolerable dose denoted by *c*_*L*_ and *c*_*U*_ respectively. The therapeutic window is shown in green. (a) and (b) drug concentration with the smallest hazard is the minimum effective dose. (c) and (d) drug concentration with the smallest hazard is the maximum tolerable dose.

## A Specific Example

To illustrate the general theory we now consider an explicit model for the within-host dynamics of infection and resistance. We model an acute infection in which the pathogen elicits an immune response that can clear the infection. Treatment is nevertheless called for because, by reducing the pathogen load, it reduces morbidity and mortality (see Appendix C for details).

We begin by considering a situation in which the maximum tolerable drug concentration *c*_*U*_ causes significant suppression of the resistant strain (Figure 2a). We stress however that if this were true then, by definition, the resistant strain is not really HLR and thus there really is no resistance problem to begin with. We include this extreme example as a benchmark against which comparisons can be made.

**Figure 2.**
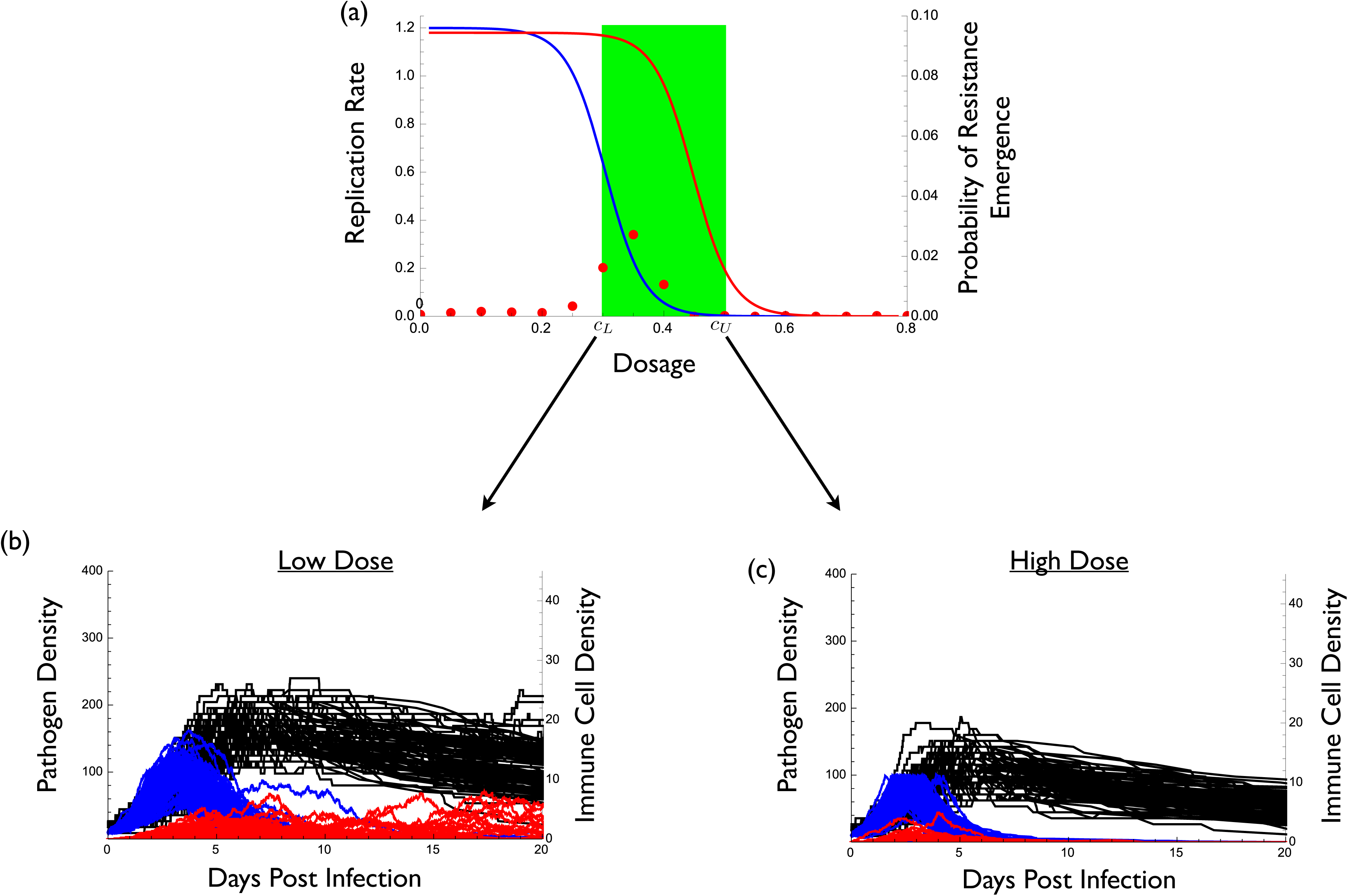
An example in which the conventional strategy of high-dose chemotherapy best prevents the emergence of resistance. (a) The dose-response curves for the wild type in blue (*r*(*c*) = 0.6(1 − tanh(15(*c* − 0.3)))) and the resistant strain in red (*r*_*m*_(*c*) = 0.59(1 − tanh(15(*c* − 0.45)))) as well as the therapeutic window in green. Red dots indicate the probability of resistance emergence. Probability of resistance emergence is defined as the fraction of 5000 simulations for which resistance reached a density of at least 100 (and thus caused disease).(b) and (c) wild type density (blue), resistant density (red), and immune molecule density (black) during infection for 1000 representative realizations of a stochastic implementation of the model. (b) treatment at the smallest effective dose *c*_*L*_, (c) treatment at the maximum tolerable dose *c*_*U*_. Parameter values are *P*(0) = 10, *P*_*m*_(0) = 0, *I*(0) = 2, *α* = 0.05, *δ* = 0.05, *κ* = 0.075, *μ* = 10^−2^, and *γ* = 0.01.

Not surprisingly, under these conditions a large dose is most effective at preventing resistance (compare Figure 2b with 2c). This is a situation in which the conventional ‘hit hard’ strategy is best.

Now suppose that the maximum tolerable drug concentration *c*_*U*_ is not sufficient to directly suppress the resistant strain (Figure 3a). In this case the only difference from Figure 2 is a change in the resistant strain’s dose-response curve. Now there really is a potential resistance problem in the sense that, from a clinical standpoint, the drug is largely ineffective against the resistant strain.

**Figure 3.**
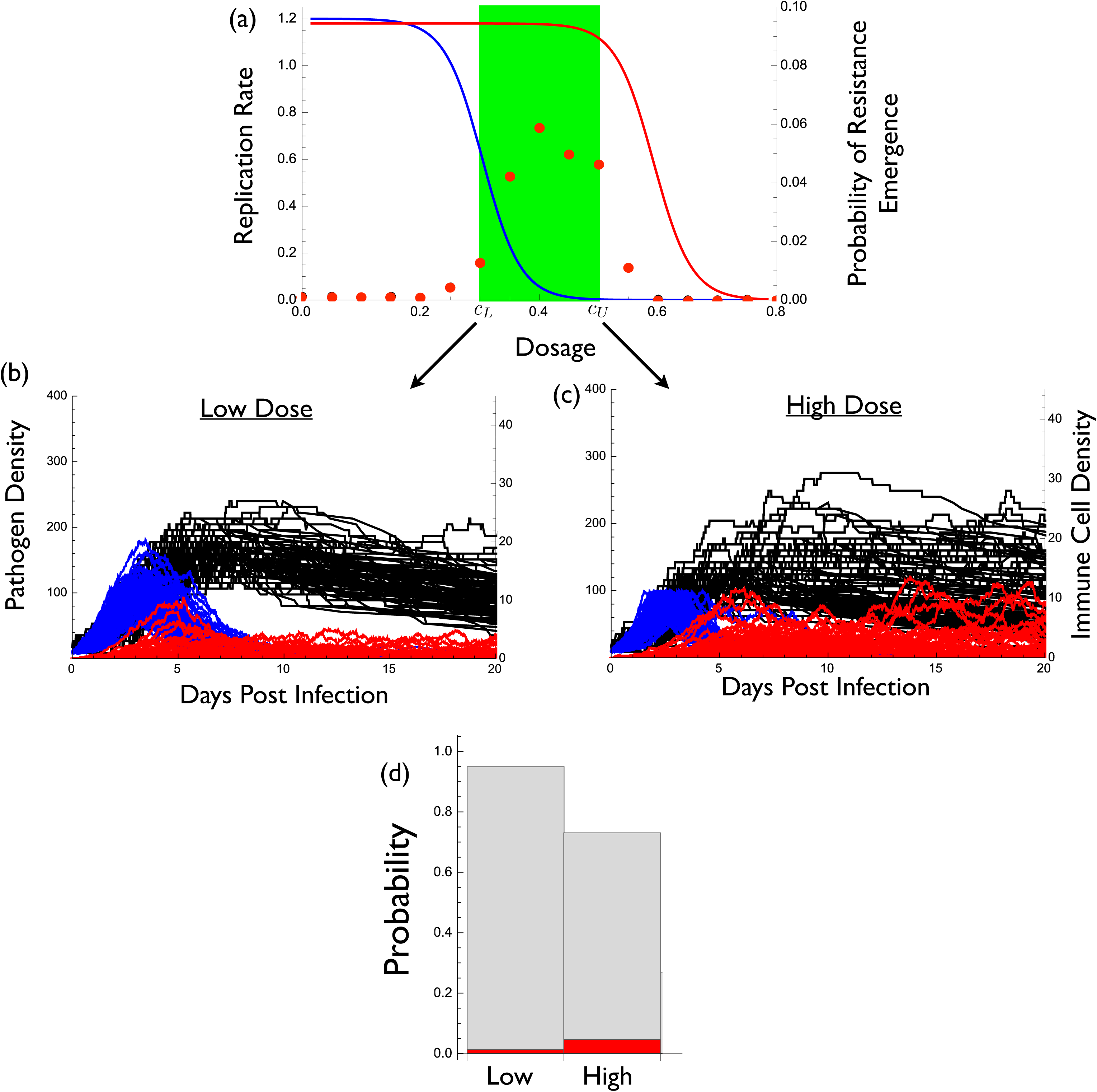
An example in which the low-dose strategy best prevents the emergence of resistance. (a) The dose-response curves for the wild type in blue (*r*(*c*) = 0.6(1 − tanh(15(*c* − 0.3)))) and the resistant strain in red (*r*_*m*_(*c*) = 0.59(1 − tanh(15(*c* − 0.6)))) as well as the therapeutic window in green. Red dots indicate the probability of resistance emergence. Probability of resistance emergence is defined as the fraction of 5000 simulations for which resistance reached a density of at least 100 (and thus caused disease). (b) and (c) wild type density (blue), resistant density (red), and immune molecule density (black) during infection for 1000 representative realizations of a stochastic implementation of the model. (b) treatment at the smallest effective dose *c*_*L*_, (c) treatment at the maximum tolerable dose *c*_*U*_. The probability that a resistant strain appears by mutation is indicated by grey bars for low and high dose. The probability of resistance emergence is indicated by the height of the red bars for these cases. The probability of resistance emergence, given a resistant strain appeared by mutation, can be interpreted as the ratio of the red to grey bars. Parameter values are *P*(0) = 10, *P*_*m*_(0) = 0, *I*(0) = 2, *α* = 0.05, *δ* = 0.05, *κ* = 0.075, *μ* = 10^−2^, and *γ* = 0.01.

Under these conditions we see that a small dose is more effective at preventing resistance emergence than a large dose (compare Figure 3b with 3c). This is a situation in which the conventional or orthodox ‘hit hard’ strategy is not optimal.

Equation (4) provides insight into these contrasting results. The only difference between the models underlying Figures 2 and 3 is that *∂π/∂c* and *∂π*^0^*/∂c* are both negative for Figure 2 whereas they are nearly zero for Figure 3 (that is, at tolerable doses, the drug has negligible effects on resistant mutants). As a result, the negative terms in equation (4) outweigh the positive terms for Figure 2 whereas the opposite is true for Figure 3.

These results appear to contradict those of a recent study by Ankomah and Levin (12). Although their model is more complex than that used here, equation (4) and its extensions in the appendices show that such additional complexity does not affect our qualitative conclusions. Ankomah and Levin (12) defined resistance evolution in two different ways: (i) the probability of emergence, and (ii) the time to clearance of infection. For the sake of comparison, here we focus on the probability of emergence. Ankomah and Levin (12) defined emergence as the appearance of a single resistant microbe. As such their emergence is really a measure of the occurrence of resistance mutations rather than emergence *per se*.

In comparison, we consider emergence to have occurred only once the resistant strain reaches clinically significant levels; namely, a density high enough to cause symptoms or to be transmitted. There are two process that must occur for *de novo* resistant strains to reach clinically relevant densities. First, the resistant strain must appear by mutation, and both our results (Figure 3d) and those of Ankomah and Levin (12) show that a high dose better reduces the probability that resistance mutations occur (this can also be seen in equation 4). Second, the resistant strain must replicate to clinically significant levels. Ankomah and Levin (12) did not account for this effect and our results show that a high concentration is worse for controlling the replication of resistant microbes *given a resistant strain has appeared* (Figure 3d). This is because higher doses maximally reduce competitive suppression. In Figure 3 the latter effect overwhelms the former, making low-dose treatment better. In Figure 2 these opposing processes are also acting but in that case the drug’s effect on controlling mutation outweighs its effect on increasing the replication of such mutants once they appear.

More generally, Figure 4 illustrates the relationship between drug concentration and the maximum size of the resistant population during treatment, for the model underlying Figure 3. In this example a high concentration tends to result in relatively few outbreaks of the resistant strain but when they occur they are very large. Conversely, a low concentration tends to result in a greater number of outbreaks of the resistant strain but when they occur they are usually too small to be clinically significant.

**Figure 4.**
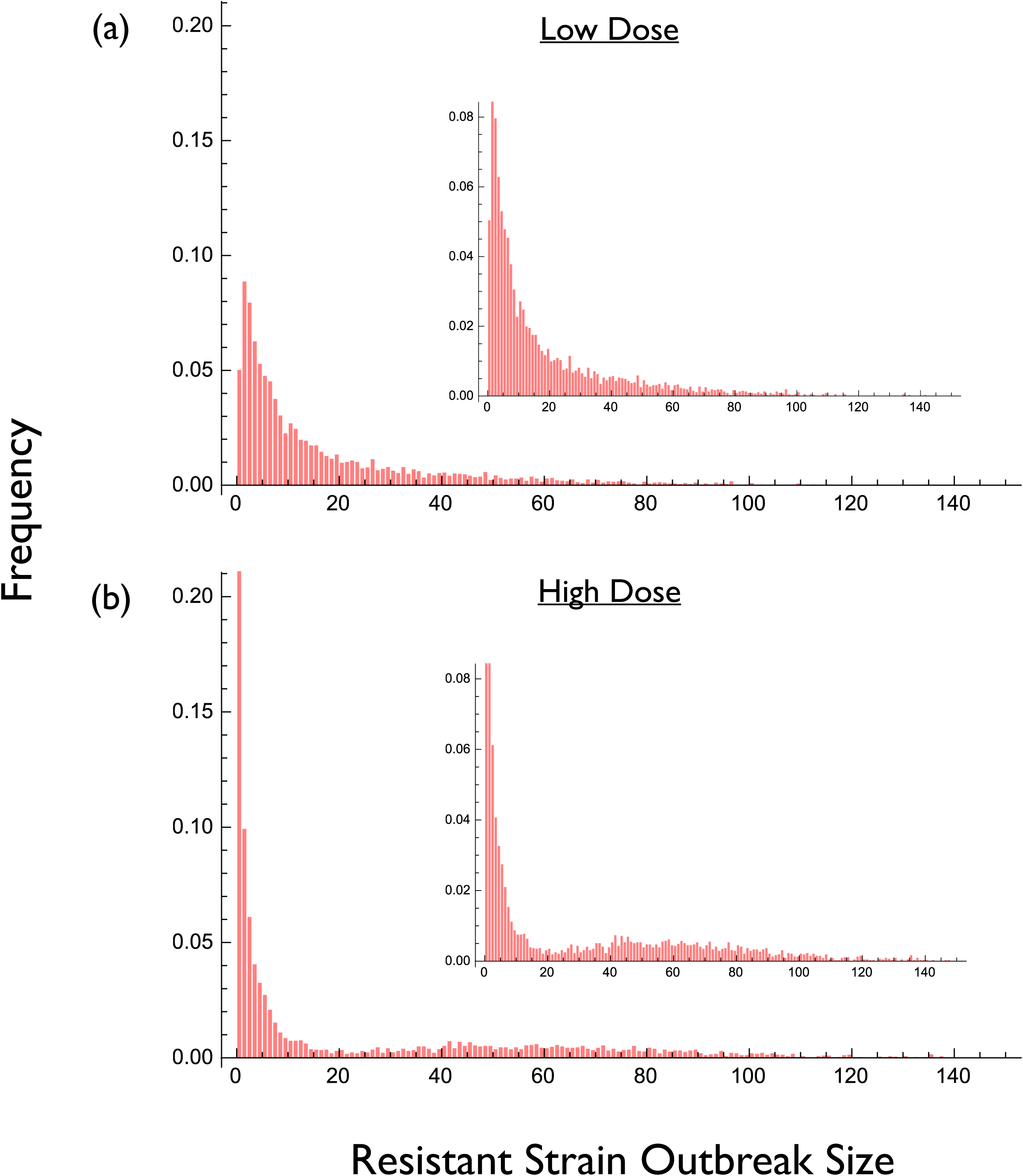
Frequency distribution of resistant strain outbreak sizes for the simulation underlying Figure 3. Each distribution is based on 5000 realizations of a stochastic implementation of the model. (a) Low drug dose. (b) High drug dose. Insets show the same distribution on a different vertical scale.

One can also examine other metrics like duration of infection, total resistant strain load during treatment, likelihood of resistant strain transmission, etc. but the above results are sufficient to illustrate that no single, general, result emerges. Whether a high or low dose is best for managing resistance will depend on the specific context (i.e., the parameter values) as well as the metric used for quantifying resistance emergence. In Appendices C and D, we consider cases where there is pre-existing resistance at the start of infection, strains with intermediate resistance, and other measures of drug dosing and resistance emergence. None of these factors alters the general finding that the optimal strategy depends on the balance between competing evolutionary processes.

## Discussion

Equation (4) clearly reveals how high-dose chemotherapy gives rise opposing evolutionary processes in the emergence of resistance. It shows how the balance between mutation and competition determines the optimal resistance management strategy (13, 19). Increasing the drug concentration reduces mutational inputs into the system but it also unavoidably reduces the ecological control of any HLR pathogens that are present. These opposing forces generate an evolutionary hazard curve that is unimodal. Consequently, the worst approach is to treat with intermediate doses (Figure 1) as many authors have recognized (5–7, 9). The best approach is to administer either the largest tolerable dose or the smallest clinically effective dose (that is, the concentration at either end of the therapeutic window). Which of these is optimal depends on the relative positions of the hazard curve and the therapeutic window (Figure 1). Administering the highest tolerable dose can be a good strategy (Figure 1c,d) but it can also be less than optimal (Figure 1b) or even the worst thing to do (Figure 1a). Thus, nothing in evolutionary theory supports the contention that a ‘hit-hard’ strategy is a good rule of thumb for resistance management.

### Empirical evidence

Our framework makes a number of empirical predictions that are consistent with existing data. First, the resistance hazard will be a unimodal function of drug concentration. This is well-verified in numerous studies. In fact a unimodal relationship between resistance emergence and drug concentration (often called an ‘inverted-U’ in the literature) is arguably the single-most robust finding in all of the empirical literature (e.g., 22–40).

Second, the position and shape of the hazard curve will vary widely among drugs and microbes, depending on how drug dose affects mutation rates and the strength of competition. Such wide variation is seen (e.g., 22, 23, 27, 28, 34, 37, 38, 41, 42), presumably reflecting variation in the strength of the opposing processes highlighted by equation (4).

Third, the relationship found between drug concentration and resistance evolution in any empirical study will depend on the range of concentrations explored. At the low end, increasing dose should increase resistance evolution; at the high end, increasing dose should decrease resistance evolution. Examples of both cases are readily seen, often even within the same study (e.g., 15, 22–40, 43–49). It is important to note that there are clear examples for which low-dose treatments can better prevent resistance emergence than high doses (15, 38, 41, 43–46, 48–50), despite an inherent focus in the literature on experimental exploration of high-dose chemotherapy. The theory presented here argues that uniformity is not expected and the bulk of the empirical literature is consistent with this prediction.

### Theory does not support using the MPC as a rule of thumb

An important and influential codification of Ehrlich’s ‘hit hard’ philosophy is the concept of the mutant selection window, and the idea that there exists a mutant prevention concentration (MPC) that best prevents resistance evolution (7–9). The MPC is defined as ‘the lowest antibiotic concentration that prevents replication of the least susceptible single-step mutant’ (see 8, p. S132). When drug concentrations are maintained above the MPC, ‘pathogens populations are forced to acquire two concurrent resistance mutations for replication under antimicrobial therapy’ (see 51, p. 731). Below the MPC lies the ‘mutant selection window’, where single-step resistant mutants can replicate, thus increasing the probability that microbes with two or more resistance mutations will appear. Considerable effort has been put into estimating the MPC for a variety of drugs and microbes (4).

The relationship between these ideas and the theory presented here is best seen using the extension of equation (4) that allows for strains with intermediate resistance. Appendix B shows that, in this case, equation (4) remains unchanged except that its first term (the mutational component) is extended to account for all of the ways in which the HLR strain can arise by mutation through strains with intermediate resistance (see expression B-3 in Appendix B). A focus on the MPC can therefore be viewed as a focus on trying to control only the mutational component of resistance emergence. And as the theory embodied by equation (4) shows, doing so ignores the other evolutionary process of competitive release that is operating. The use of the MPC therefore cannot be supported by evolutionary theory as a general rule of thumb for resistance management.

If evolutionary theory does not support the use of MPC as a general approach then why does this nevertheless appear to work in some cases (e.g., 33, 52)? The theory presented here provides some possible explanations. First, if HLR strains can appear only through mutation from strains with intermediate resistance, and if feasible dosing regimens can effectively kill all first step mutants, then such an approach must necessarily work since it reduces all mutational input to zero. But for most of the challenging situations in medicine, achieving this is presumably not possible. For example, if the MPC is not delivered to all pathogens in a population because of patient compliance, metabolic variation, spatial heterogeneity in concentration, etc, then the mutational input will not be zero. Also, if HLR strains can arise in ways that do not require mutating through strains with intermediate resistance (e.g., through lateral gene transfer; 53) then again the mutational input will not be zero. In either case, one must then necessarily account for how the choice of dose affects the opposing evolutionary process of competitive release in order to minimize the emergence of resistance. Figure C3 in Appendix C illustrates this idea by presenting a numerical example in which the MPC is the worst choice of drug concentration for controlling HLR.

Second, the theory presented here suggests that the MPC *can* be the best way to contain resistance if this concentration happens to be the upper bound of the therapeutic window (although see Figure C3 of Appendix C for a counterexample). If, however, the MPC is less than the upper bound then even better evolution-proofing should be possible at either end of the therapeutic window. If the MPC is greater than the upper bound, as it is for example with most individual TB drugs (54) and levofloxacin against *S. aureus* (27), the MPC philosophy is that the drug should then be abandoned as monotherapy. But our framework suggests that before doing so, it might be worthwhile considering the lower bound of the therapeutic window. Researchers have tended not to examine the impact of the smallest clinically effective dose on resistance evolution, perhaps because of an inherent tendency to focus on high-dose chemotherapy. It would be informative to compare the effects of the MPC with concentrations from both ends of the therapeutic window on resistance emergence experimentally.

### Theory does not support using the highest tolerable dose as a rule of thumb

The MPC has yet to be estimated for many drug-microbe combinations (4) and it can be difficult to do so, especially in a clinically-relevant setting (51, 53). Given the uncertainties involved, and the need to make clinical decisions ahead of the relevant research, some authors have suggested the working rule of thumb of administering the highest tolerable dose (3, 4). Our analysis shows that evolutionary theory provides no reason to expect that this approach is best. By reducing or eliminating the only force which retards the emergence of any HLR strains that are present (i.e., competition), equation (4) makes clear that a hit hard strategy can backfire, promoting the very resistance it is intended to contain.

### How to choose dose

If the relative positions of the HLR hazard curve and the therapeutic window are known, rational (evidence-based) choice of dose is possible. If the therapeutic window includes doses where the resistance hazard is zero, then those doses should be used. However, by definition, such situations are incapable of generating the HLR which causes a drug to be abandoned, and so these are not the situations that are most worrisome. If the hazard is non-zero at both ends of the therapeutic window, the bound associated with the lowest hazard should be used (Figure 1b, c). If nothing is known of the HLR hazard curve (as will often be the case), then there is no need to estimate the whole function. Our analysis suggests that the hazards need be estimated only at the bounds of the therapeutic window. These bounds are typically well known because they are needed to guide clinical practice. Estimating the resistance hazard experimentally can be done in vitro and in animal models but we note that since the solution falls at one end of the therapeutic window, they can also be done practically and ethically in patients. That will be an important arena for testing, not least because an important possibility is that, as conditions change, the optimal dose might change discontinuously from the lowest effective dose to the highest tolerable dose or vice versa. There is considerable scope to use mathematical and animal models to determine when that might be the case and to determine clinical predictors of when switches should be made.

### Managing resistance in non-targets

Our focus has been on the evolution of resistance in the pathogen population responsible for disease. Looking forward, an important empirical challenge is to consider the impact of drug dose on the broader microbiome. Resistance can also emerge in non-target micro-organisms in response to the clinical use of antimicrobials (44). Resistance in those populations can increase the likelihood of resistance in future pathogen populations, either because of lateral gene transfer from commensals to pathogens, or when commensals become opportunistic pathogens (9, 55). For instance, aggressive drug treatment targeted at bacterial pneumonia in a rat model selected for resistance in gut fauna. Lower dose treatment of the targeted lung bacteria was just as clinically effective and better managed resistance emergence in the microbiota (50).

It is unclear just how important these off-target evolutionary pressures are for patient health, but if they are quantitatively important, this raises the interesting and challenging possibility that the real hazard curve should be that of the collective microbiome as a whole, weighted by the relative risk of resistance evolution in the components of the microbiome and the target pathogen. It will be challenging to determine that, but our focus on either end of the therapeutic window at least reduces the parameter space in need of exploration.

## Coda

Our analysis suggests that resistance management is best achieved by using a drug concentration from one edge of the therapeutic window. In practice, patients are likely treated somewhat more aggressively than the minimum therapeutic dose (to ensure no patients fail treatment) and somewhat less aggressively than the maximum tolerable dose (to ensure no patients suffer toxicity). This means that medical caution is always driving resistance evolution faster than it need go, particularly when the maximum hazard lies within the therapeutic window (Figure 1b,c). From the resistance management perspective, it is important to determine the level of caution that is clinically warranted rather than simply perceived.

For many years, physicians have been reluctant to shorten antimicrobial courses, using long courses on the grounds that it is better to be safe than sorry. It is now increasingly clear from randomized trials that short courses do just as well in many cases (e.g., 56–58) and they can reduce the risk of resistance emergence (56, 59, 60). We suggest that analogous experiments looking at the evolutionary outcomes of lowest clinically useful doses should be the next step. Such experiments in plants have already shown unambiguously that low dose fungicide treatment best prevents the spread of resistant fungal pathogens (61). How generally true this is for other pathogens, or pathogens of other hosts, remains to be seen. We also note that our arguments about the evolutionary merits of considering the lowest clinically useful doses have potential relevance in the evolution of resistance to cancer chemotherapy as well (62).

## Acknowledgements

We thank the Research and Policy in Infectious Disease Dynamics (RAPIDD) program of the Science and Technology Directorate, the Department of Homeland Security, the Fogarty International Center, and the National Institutes of Health for support. We also thank R. Woods, E. Hansen, and members of the Princeton RAPIDD workshop organized by Metcalf and R. Kouyos for discussion. This research was also supported by a grant from the Natural Sciences and Engineering Research Council of Canada (TD) and by the National Institute of General Medical Sciences (R01GM089932) (AFR). The funders had no role in study design, data collection and analysis, decision to publish, or preparation of the manuscript.

## APPENDICES A-E

## Appendix A - Derivation of Equation 4

In the absence of treatment we model the within-host dynamics using a system of differential equations

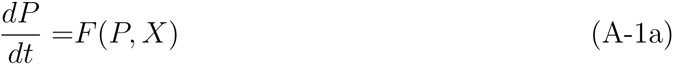

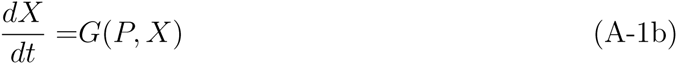

where *P* is the density of the wild type and *X* is a vector of variables describing the within-host state (e.g., RBC count, densities of different immune molecules, etc). The initial conditions are *P*(0) = *P*_0_ *X*(0) = *X*_0_. At some point, *t*^*\**^, drug treatment is introduced. Using lower case letters to denote the dynamics in the presence of treatment, we then have

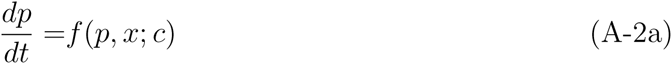

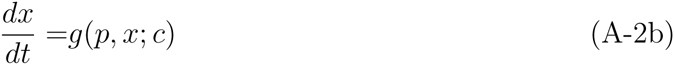

with initial conditions *p*(0; *c*) = *P*(*t*^*\**^) and *x*(0; *c*) = *X*(*t*^*\**^), and where *c* is the dosage. For simplicity, here we assume that a constant drug concentration is maintained over the course of the infection. Appendix E considers the pharmacokinetics of discrete drug dosing. The notation *p*(*t*; *c*) and *x*(*t*; *c*) reflects the fact that the dynamics of the wild type and the host state will depend on dosage. For example, if the dosage is very high *p* will be driven to zero very quickly.

As the drug removes the wild type pathogen, resistant mutations will continue to arise from the wild type population stochastically. For example, if mutations are produced only during replication of the wild type, then the rate of mutation will have the form *μr*(*c*)*p*(*t*; *c*) where *μ* is the mutation rate and *r*(*c*) is the replication rate of the wild type pathogen (which depends on drug dosage *c*). With this form of mutation, if we could administer the drug at concentrations above the MIC at the very onset of infection, then resistance evolution through *de novo* mutation would not occur. In reality symptoms and therefore drug treatment typically do not occur until later in the infection, meaning that some resistant strains might already be present at low frequency at the onset of treatment. There are also other plausible forms for the mutation rate as well, and therefore we simply specify this rate by some general function *λ*[*p*(*t*; *c*), *c*].

Whenever a resistant strain appears it is subject to stochastic loss. We define *π* as the probability of avoiding loss (which we refer to as ‘escape’). To simplify the present analysis, we use a separation of timescales argument and assume that the fate of each mutant is determined quickly (essentially instantaneously) relative to the dynamics of the wild type and host state (we relax this assumption in all numerical examples). Thus, *π* for any mutant will depend on the host state at the time of its appearance, *x*(*t*; *c*), and it will therefore depend indirectly on *c*. Note that *π* will also depend directly on *c*, however, because drug dosage might directly suppress resistant strains as well if the dose is high enough. Therefore we use the notation *π*[*x*(*t*; *c*), *c*], and assume that *π* is an increasing function of *x* and a decreasing function of *c*.

With the above assumptions the host can be viewed as being in one of two possible states at any point in time during the infection: (i) resistance has emerged (i.e., a resistant strain has appeared and escaped), or (ii) resistance has not emerged. We model emergence as an inhomogeneous birth process, and define *q*(*t*) as the probability that resistance has emerged by time *t*. A conditioning argument gives

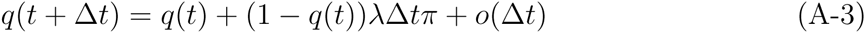

where *λ*Δ*t* is the probability that a mutant arises in time Δ*t*, and *π* is the probability that such a mutant escapes. Re-arranging and taking the limit Δ*t →* 0 we obtain

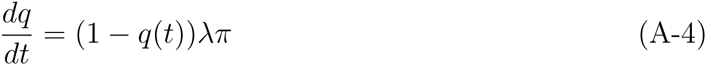

with initial condition *q*(0) = *q*_0_. Note that *q*_0_ is the probability that emergence occurs as a result of resistant mutants being present at the start of treatment. Again employing a separation of timescales argument, if there are *n* mutant individuals present at this time, then *q*_0_ = 1 − (1 − *π*[*x*(0; *c*), *c*])^*n*^.

The solution to the above differential equation is

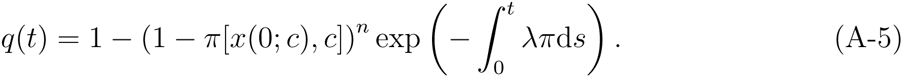

If *a* is the time at which treatment is stopped, and *Q* is the probability of emergence occurring at some point during treatment, then *Q* = *q*(*a*). If we further define *S* = −*n* ln (1 − *π*[*x*(0; *c*), *c*]) then we can write *Q* as

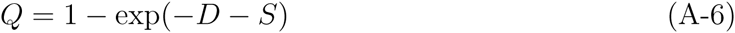

where 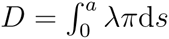. We refer to *D* as the *de novo* hazard and *S* as the standing hazard. *D* is the contribution to escape that is made up of mutant individuals that arise during the course of treatment. *S* is the contribution to escape that is made up of mutant individuals already presents at the start of treatment.

Given the expression for *Q*, all else equal, resistance management would seek the treatment strategy, *c* that makes *Q* as small as possible. Since *Q* is a monotonic function of *D* + *S*, we can simplify matters by focusing on these hazards instead. Thus we define

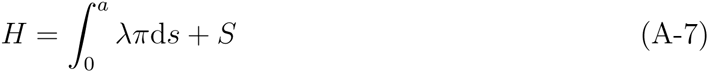

which is the ‘total hazard’ during treatment. Equation (4) is then obtained by differentiating the the total hazard *H* with respect to *c*.

## Appendix B - Extensions involving intermediate strains and horizontal gene transfer

The results of the main text (which are derived in Appendix A) are based on the assumption that a single mutational event can give rise to high-level resistance. In some situations several mutational events might be required. These so-called ‘stepping stone mutations’ towards high-level resistance might themselves confer an intermediate level of resistance. One of the arguments in favour of aggressive chemotherapy has been to prevent the persistence of these stepping stone strains, and thereby better prevent the emergence of high-level resistance (1–8). Here we incorporate such stepping stone mutations into the theory, again placing primarily attention on the emergence of high-level resistance.

As in Appendix A, in the absence of treatment we model the within-host dynamics using a system of differential equations

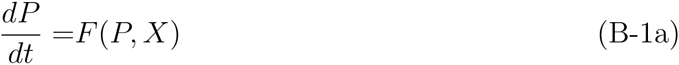

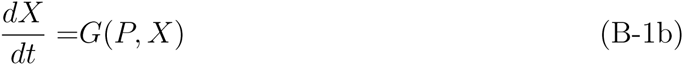

but now *P* is also a vector containing the density of the wild type and all potential intermediate mutants. All intermediate strains are assumed to bear some metabolic or replicative cost as well, meaning that they are unable to increase in density in the presence of the wild type. Mechanistically again this is because the wild type has suppressed the host state, *X*, below the minimum value required for a net positive growth by any intermediate strain. Thus, in the absence of treatment we expect most of these mutants to have negligible density. Once treatment is introduced we have

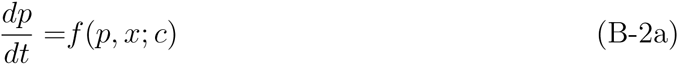

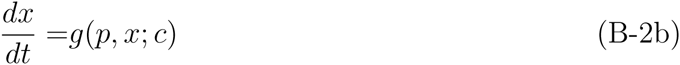

where again *p* is now a vector. As before we have initial conditions *p*(0; *c*) = *P*(*t*^*\**^) and *x*(0; *c*) = *X*(*t*^*\**^), and where *c* is the dosage. Now, however, different choices of *c* will generate different distributions of strain types *p*(*t*; *c*) during the infection. Furthermore, each type will give rise to the high-level resistance strain with its own rate. Therefore, the function specifying the rate of mutation to the HLR strain *λ*[*p*(*t*; *c*), *c*] is a function of the vector variable *p*(*t*; *c*).

The calculations in Appendix A can again be followed. We obtain an equation identical to equation (4) except that the first term is replaced by

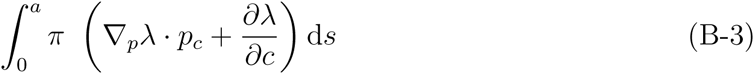

where subscripts denote differentiation with respect to that variable. The difference is that (*∂λ/∂p*)(*∂p/∂c*) in equation (4) is replaced with ∇_*p*_*λ · p*_*c*_. The quantity *p*_*c*_ is a vector whose components are the changes in the density of each intermediate strain arising from an increased dosage. The quantity ∇_*p*_*λ* is the gradient of the mutation rate with respect to a change in the density of each intermediate strain. The integral of the dot product of the two, ∇_*p*_*λ · p*_*c*_, is therefore the overall change in mutation towards the HLR strain during treatment. Whereas the first term of equation (4) is expected to be negative, expression (B-3) can be negative or positive depending on how different doses affect the distribution of intermediate mutants during the infection (i.e., the elements of *p*_*c*_) and the rate at which each type of intermediate mutant gives rise to the strain with high level resistance (i.e., the elements of ∇_*p*_*λ*). Either way, however, this does not alter the salient conclusion that the optimal resistance management dose will depend on the details.

In an analogous fashion we might also alter the derivation in Appendix A to account for the possibility that some microbes acquire high-level resistance via horizontal gene transfer from other, potentially commensal, microbes. To do so we would simply need to alter the way in which *λ* is modelled. In particular, it might then be a function of the densities of commensal microbes as well, who themselves could be affected by drug dosage. Thus, once treatment has begun, we might have a system of equations of the form

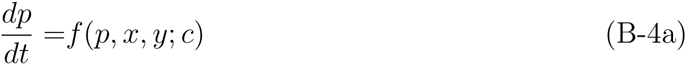

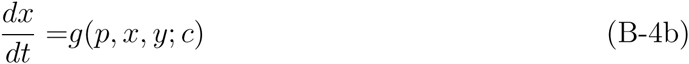

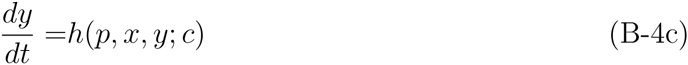

where *y* is a vector of commensal microbe densities. We might then model *λ* as *λ*[*p*(*t*; *c*), *y*(*t*; *c*)]. Again, calculations analogous to those of Appendix A can be followed to obtain an appropriate expression for the resistance hazard. As with the above examples, there will again be a tradeoff between components of this expression as a function of drug dosage.

## Appendix C - A Model of acute immune-mediated infections

The dynamics of the mutant and wild type in the absence of treatment are modeled as

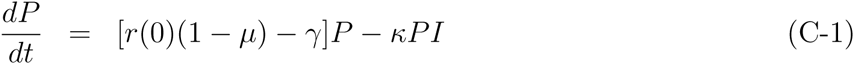

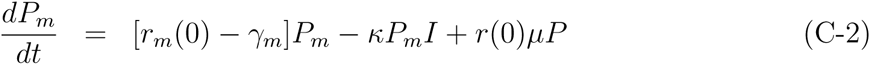

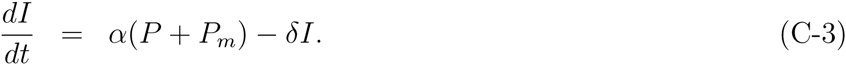

where *r*(·) and *r*_*m*_(·) are the growth rates of the wild type and mutant as a function of drug concentration, *μ* is the mutation probability from wild type to resistant, and *γ* and *γ*_*m*_ are the natural death rates of each. We assume a cost of resistance in the absence of treatment, meaning that *r*(1 − *μ*) − *γ > r*_*m*_ − *γ*_*m*_ The immune response, *I*, grows in proportion to the density of the pathogen population and decays at a constant per capita rate *δ*. Immune molecules kill the pathogen according to a law of mass action with parameter *κ* for both the wild type and the resistant strain (i.e., immunity is completely cross-reactive). This is a simple deterministic model for an immune-controlled infection. When the mutation rate is zero (*μ* = 0) and the pathogen can increase when rare, the model displays damped oscillations towards an equilibrium with the wild type present 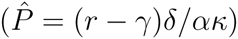, the mutant extinct 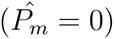, and the immune system at a nonzero level 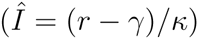. For many choices of parameter values (including those that we focus on here) the first trough in pathogen density is very low, and therefore once we introduce stochasticity the entire pathogen population typically goes extinct at this stage, at which point the immune molecules then decay to zero. It is in this way that we model an immune-controlled infection.

Under treatment the dynamics are the same as above but where *r*(·) and *r*_*m*_(·) are then evaluated at some nonzero drug concentration. Throughout we assume that the dose-response functions *r*(*·*) and *r*_*m*_(*·*) are given by the function *b*_1_(1 − tanh(*b*_2_(*c − b*_3_))) for some constants *b*_1_, *b*_2_, and *b*_3_. The model used to explore the emergence of resistance employs a stochastic implementation of the above equations using the Gillespie algorithm.

Figure C1 presents output for several runs of the model using three different drug concentrations. In all cases we have set the mutation rate to zero (no resistant strains arise). In the absence of treatment an infection typically results in a single-peak of wild type pathogen before the infection is cleared. To model realistic disease scenarios we (arbitrarily) suppose that infected individuals become symptomatic only once the pathogen density exceeds a threshold of 100 and treatment is used only once an infection is symptomatic. For the parameter values chosen in this example, 99% of untreated infections are symptomatic (Figure C1a,b). We further suppose (again arbitrarily) that a pathogen load greater than 200 results in substantial morbidity and/or mortality. With this assumptions we can then proceed to define the therapeutic window. The upper limit *c*_*U*_ is arbitrary in the model and so we set *c*_*U*_ = 0.5. The lower limit *c*_*L*_ is the smallest dose that prevents significant morbidity and/or mortality. Therefore it is the smallest dose that, in the absence of resistance emergence, keeps pathogen load below 200. Figure C1c shows that, for the parameter values used, *c*_*L*_ *≈* 0.3. Notice from Figure C1a that a dose of *c* = 0.3 does not fully suppress growth as measured *in vitro* but it nevertheless controls the infection *in vivo* because the immune response also contributes to reducing the pathogen load.

For simulations in which the mutation rate to resistance is non-zero we quantify the emergence of resistance in the following way. For each simulation run we record the maximum density of the resistant strain before the infection is ultimately cleared. Runs in which this density reaches a level high enough to cause symptoms (a density of 100 in this case) are deemed to be infections in which resistance has ‘emerged’. The probability of resistance emergence is quantified as the fraction of runs in which this threshold level is reached. In Figure 4 of the text we also consider the consequences of using other threshold densities to define emergence.

The simulation results of the main text assume that all resistant strains arise *de novo* in a infection but in some cases we might expect resistant strains to already be present at the start of infection. The general theory presented in the main text reveals that again we should not expect any simple generalities. For example, one might expect that when the initial infection already contains many resistant microbes the relevance of *de novo* mutation might be diminished and so a lower dose might be optimal for managing resistance. Although this is sometimes the case (Day, unpubl. results) the opposite is possible as well.

As an example, Figure C2 presents results for the probability of emergence as a function of dose, for three different levels of resistance frequency in the initial infection. As the frequency of resistance in the initial infection increases, the optimal concentration changes from a low dose to a high dose. The reason is that, if resistance is already very common early in the infection, then the competitive release that occurs from removing the wild type is greatly diminished since the resistant strain will have already managed to gain a foothold before the wildtype numbers increase significantly. Put another way, the benefits of low dose therapy have decreased because the magnitude of competitive release (the blue terms in equation (4) of the main text) has decreased. Experimental results have verified this prediction; namely, that drug resistant pathogens can reach appreciable within-host densities in the absence of treatment if the initial infection contains a substantial number of these (9).

A common suggestion is that, when strains with intermediate levels of resistance are possible, aggressive chemotherapy is then optimal because anything less will allow these intermediate strains to persist and thereby give rise to HLR through mutation. We therefore conducted simulations to explore this idea. We note, however, that again the general theoretical results of Appendix B reveal that no generalities should be expected and our simulations bear this out. For example, we extended equations for the within-host dynamics to allow for a strain with intermediate resistance by using the following equations:

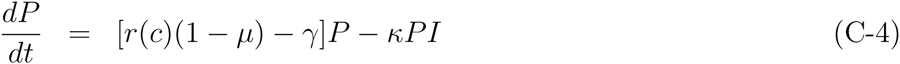

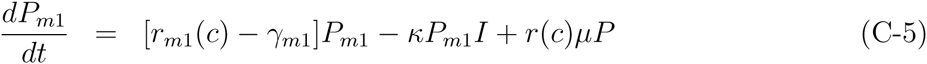

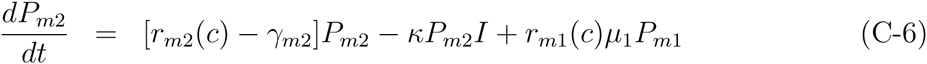

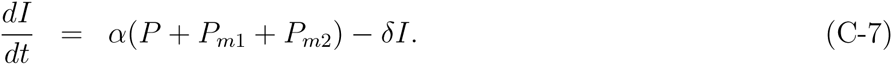

where *P*_*m*1_ is the density of the mutant strain with intermediate resistance and *P*_*m*2_ is the strain with HLR. Also, *r*(*·*), *r*_*m*1_(*·*), and *r*_*m*2_(*·*) are the growth rates of the wild type and the two mutant types as a function of drug concentration, *μ* is the mutation probability from wild type to the intermediate strain, *μ*_1_ is the mutation rate from the intermediate strain to HLR, and the *γ*’s are the natural death rates of each. Again the immune response, *I*, grows in proportion to the density of the pathogen population and decays at a constant per capita rate *δ*.

Again the simulation was conducted with a stochastic implementation of the above model using the Gillespie algorithm. While the presence of intermediate strains does alter the relative balance of factors affecting resistance emergence, this balance can still move in either direction.

As an example, Figure C3 presents simulation results in which low-dose treatment yields the lowest probability of HLR emergence. Note, however, that high-dose treatment controls the emergence of the intermediate strain the best.

The results of Figure C3 can also be interpreted within the context of the mutant selection window hypothesis and the mutant prevention concentration or MPC. The MPC is the drug concentration that prevents the emergence of all single-step resistant mutants. In Figure C3 we can see that the emergence of the intermediate, single step, mutant strain is prevented by using the maximum tolerable dose. Nevertheless, even though the HLR strain can arise only by mutation from this intermediate strain, it it the lowest effective dose that best controls the emergence of HLR. The reason for this is that it is not possible to achieve the MPC early enough in the infection to prevent all mutational input from occurring because treatment starts only once symptoms appear. For the specific case illustrated in Figure C3 the possibility of HLR arising is then enough to tip the balance so that the lower edge of the therapeutic window is the best strategy for controlling HLR.

## Appendix D - Other results for the model of acute immune-mediated infections

In the main text we focus on the emergence of the resistant strain but in many clinical studies researchers focus instead on successful treatment. For example, one common approach is to quantify the probability of treatment failure as a function of drug dose (or some proxy thereof). Such studies cannot provide information about resistance evolution *per se* but they nevertheless might involve a component of resistance evolution if this is one of the potential reasons for treatment failure.

We can explore a similar idea in the context of the model in the main text. Suppose we measure clinical success as the complete eradication of infection by day 20. In the simulations some individuals then display treatment failure because, through the stochasticity of individual infection dynamics, they fail to clear the infection by this time. Figure D1a presents the probability of treatment failure, measured by the fraction of the simulations for which the infection (wild type or resistant) was still present on day 20 for the model underlying Figure 3. Failure occurs under both treatment scenarios but it happens more frequently for the high dose treatment (compare red portion of bar graphs in Figure D1a). There is an important structure to these failures, however, that can be better appreciated by calculating the probability of failure by conditioning on whether or not a resistant mutation ever appeared during treatment; i.e.,

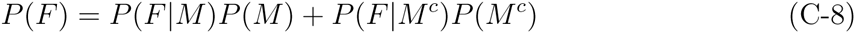

where *P*(*F*) is the probability of failure, *P*(*M*) is the probability of a resistant mutation appearing during treatment (*P*(*M* ^*c*^) is the probability that this doesn’t occur), and *P*(*F |M*) is the probability of failure given a resistant mutation appears (with *P*(*F |M* ^*c*^) the probability of failure given a resistant mutation does not appear). The bar graphs in Figure D1a show again that a high dose better controls the appearance of resistant mutations (i.e., *P*(*M*) is lower for the high dose treatment), but if a resistant mutation does occur, then a high dose results in a greater likelihood of treatment failure (i.e., *P*(*F |M*) is higher for the high dose treatment - note that this quantity can be interpreted graphically as the ratio of the red to grey bars). And in this case the latter effect overwhelms the former, making the probability of treatment failure *P*(*F*) greater overall for the high dose treatment.

It is not difficult to obtain diametrically opposite results, however, with a small change in parameter values. Figure D1b show analogous results for the very same simulation, but where the probability of mutation is an order of magnitude lower. In this case we see that, even though a high dose results in a greater probability of failure if a resistant mutation appears, the effect is diminished such that, overall, the high dose results in a lower overall probability of failure. Notice also though that, even though a high dose results in a lower likelihood of treatment failure, it nevertheless still results in a higher probability of resistance emergence during treatment. The former is measured only by whether or not the infection still persists on day 20 whereas the latter is measured by whether or not a large outbreak of resistance occurs at some point during treatment. This provides an example illustrating the general idea that treatment failure cannot be taken as a proxy for resistance emergence.

## Appendix E - Generalizing the pharmacokinetics

Here we illustrate how the qualitative conclusions of the main text hold more broadly by deriving the analogue of equation (4) for quite general forms of pharmacokinetics. For simplicity we will ignore the possibility that resistant strains might be present at the start of treatment.

For the sake of illustration we suppose that the drug is administered in some arbitrary way for a period of time of length *T* and then treatment is stopped. The question we ask is, how does increasing the duration of treatment *T* affect the probability of resistance emergence? More generally we might alter other aspects of treatment like dose size, inter-dose interval, etc but our focus on *T* will be sufficient to see how one would deal with these other factors as well.

To allow for more general pharmacokinetics we must model the dynamics of drug concentration explicitly. Once treatment has begun the model becomes

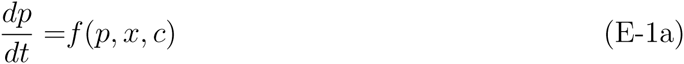

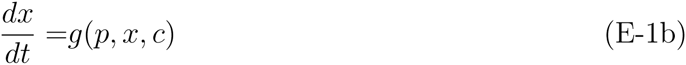

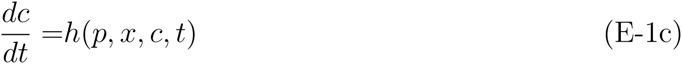

The third equation accounts for the pharmacokinetics of the drug and allows for the treatment protocol to vary through time. These equations must also be supplemented with an initial condition specifying the values of the variables at the start of treatment.

After time *T* has elapsed treatment is stopped and the dynamics then follow a different set of equations given by

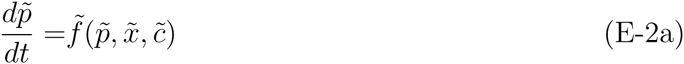

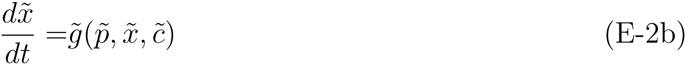

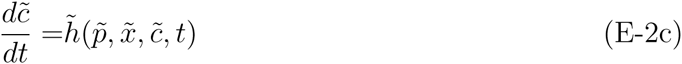

The tildes reflect the fact that the functional form of the dynamical system might change when treatment is stopped (e.g., there is no longer any input of the drug in the function 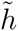 as compared with the function *h*), and thus the variables follow a different trajectory than they would have under treatment. This system of differential equation must also be supplemented with an initial condition as well, and this requires 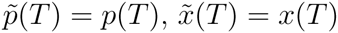, and 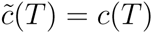. Notice that the trajectories of the new variables 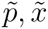 and 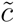 therefore depend on the duration of treatment *T* because this duration will affect their initial values. With the above formalism we can write the hazard as

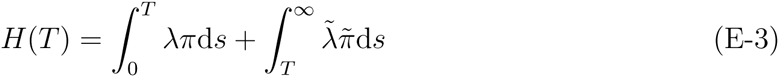

where we have simplified the notation by using a tilde above a function to indicate that the function is evaluated along the variables with a tilde. Differentiating with respect to *T* gives

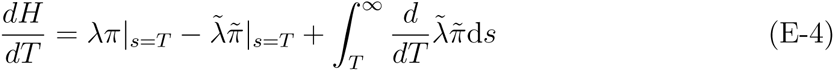

By the continuity of the state variables the first two terms cancel and therefore we have

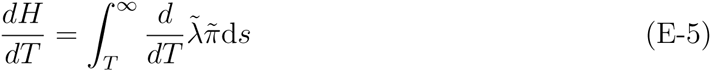

Now 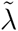 and 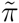 depend on *T* because they depend on the trajectories of the variables 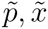 and 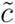 and the trajectories of these variables in turn depend on their initial conditions (which depend on *T* as described above). We can capture this notationally by treating the variables 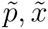 and 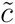 as functions of *T*. Thus we have

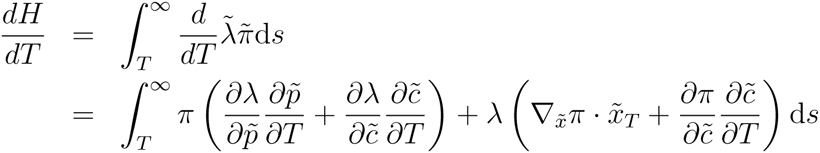

We can see that this has a form that is identical to *de novo* part of equation (4) except that now the drug concentration is no longer directly under our control. Instead, changes in *T* affect resistance emergence by how they affect changes in drug concentration. More generally, the very same potentially opposing processes as those in equation 4 will arise regardless of how we alter the drug dosing regimen because any such alteration must ultimately be mediated through its affect on the drug concentration at each point in time during an infection.

## Figure Captions

**Figure C1.**
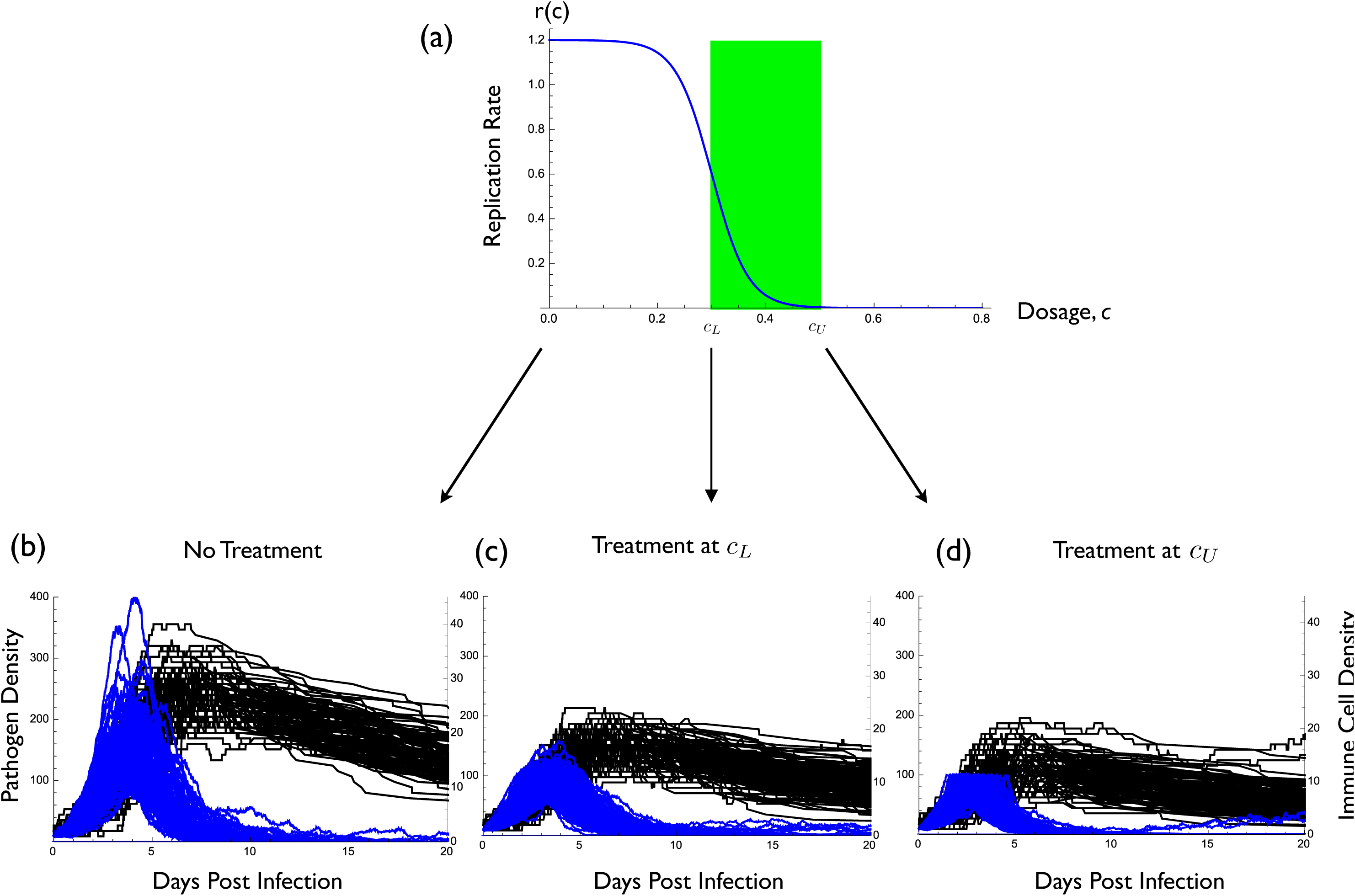
(a) The dose-response curve *r*(*c*) = 0.6(1 − tanh(15(*c* − 0.3))) as well as the therapeutic window in green. (b), (c) and (d) show wild type pathogen density (blue) and immune molecule density (black) during infection for 1000 representative realizations of a stochastic implementation of the model. (b) no treatment, (c) treatment at the smallest effective dose *c*_*L*_, (d) treatment at the maximum tolerable dose *c*_*U*_. Parameter values are *P*(0) = 10, *I*(0) = 2, *α* = 0.05, *δ* = 0.05, *κ* = 0.075, *μ* = 0, and *γ* = 0.01.

**Figure C2.**
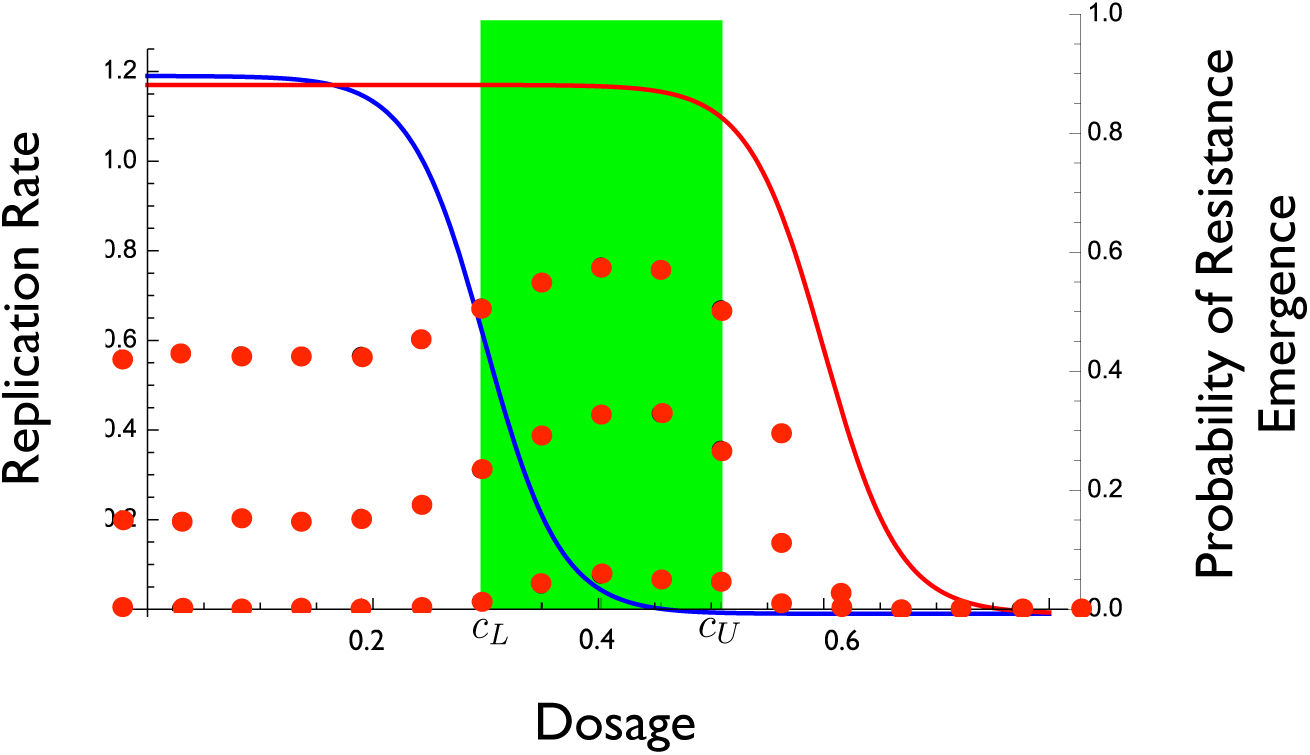
The effect of different levels of standing variation for resistance in the initial infection. Simulation is identical to that for Figure 3a except for the initial conditions. The dose-response curves for the wild type in blue (*r*(*c*) = 0.6(1 − tanh(15(*c* − 0.3)))) and the resistant strain in red (*r*_*m*_(*c*) = 0.59(1 − tanh(15(*c* −0.6)))) as well as the therapeutic window in green. Red dots indicate the probability of resistance emergence, and for three different initial conditions. Probability of resistance emergence is defined as the fraction of 5000 simulations for which resistance reached a density of at least 100 (and thus caused disease). Top set of dots have *P*(0) = 5, *P*_*m*_(0) = 5; middle set of dots have *P*(0) = 7, *P*_*m*_(0) = 3; bottom set of dots have *P*(0) = 10, *P*_*m*_(0) = 0. Other parameter values are *I*(0) = 2, *α* = 0.05, *δ* = 0.05, *κ* = 0.075, *μ* = 10^−2^, and *γ* = 0.01.

**Figure C3.**
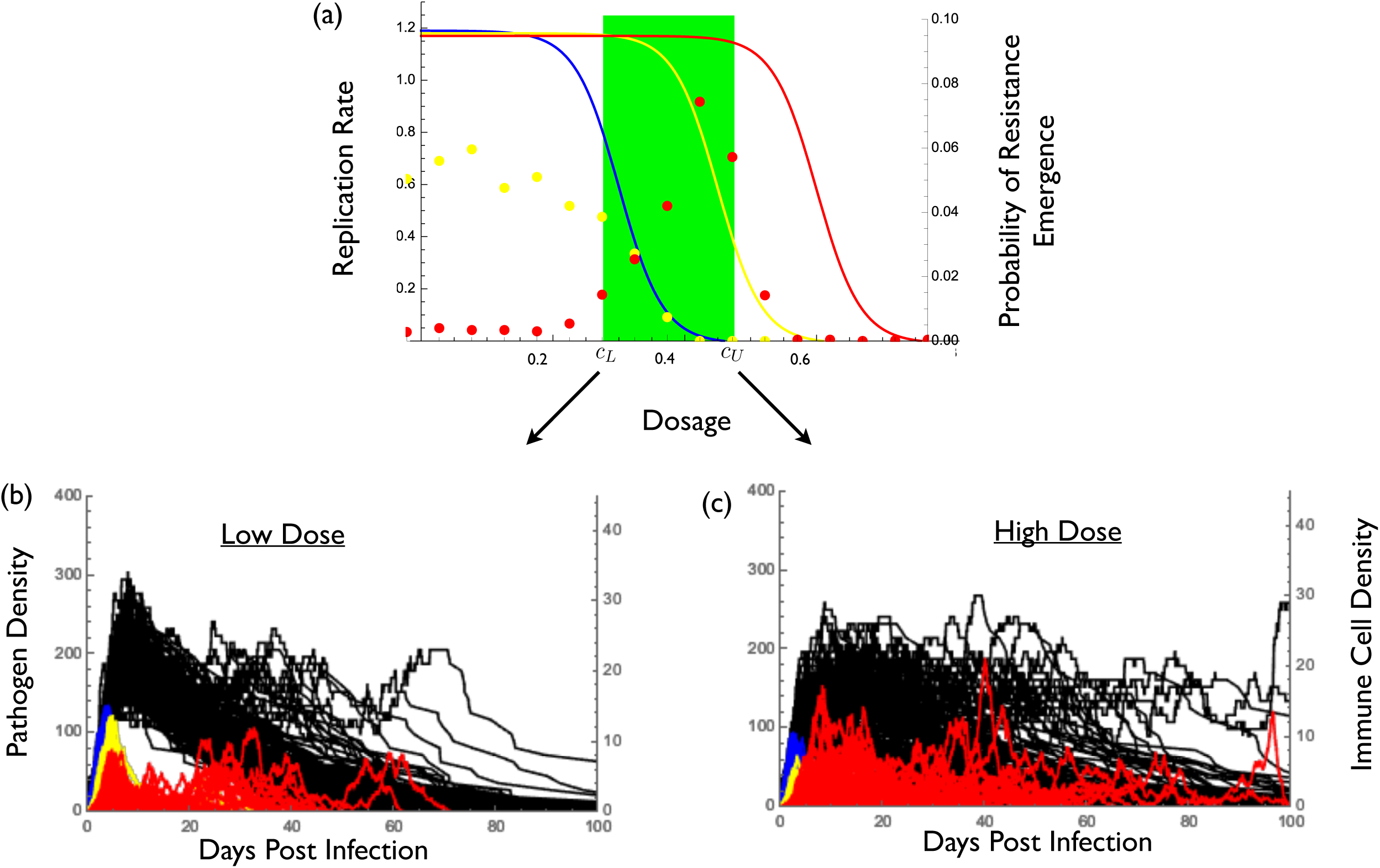
Simulation results when there is a strain with intermediate resistance. (a) The dose-response curves for the wild type in blue (*r*(*c*) = 0.6(1 − tanh(15(*c* − 0.3)))), the intermediate strain in yellow (*r*_*m*2_(*c*) = 0.595(1 − tanh(15(*c* − 0.45)))), and the HLR strain in red (*r*_*m*2_(*c*) = 0.59(1 − tanh(15(*c* − 0.6)))) as well as the therapeutic window in green. Dots indicate the probability of emergence for the intermediate strain (yellow) and the HLR strain (red). Probability of emergence is defined as the fraction of 5000 simulations for which the strain reached a density of at least 100. (b) and (c) wild type density (blue), intermediate strain density (yellow), HLR strain density (red), and immune molecule density (black) during infection for 1000 representative realizations of a stochastic implementation of the model. (b) treatment at the smallest effective dose *c*_*L*_, (c) treatment at the maximum tolerable dose *c*_*U*_. Parameter values are *P*(0) = 10, *P*_*m*1_(0) = 0, *P*_*m*2_(0) = 0, *I*(0) = 2, *α* = 0.05, *δ* = 0.05, *κ* = 0.075, *μ* = 10^−2^, *μ*_1_ = 10^−2^, and *γ* = *γ*_*m*1_ = *γ*_*m*2_ = 0.01.

**Figure D1.**
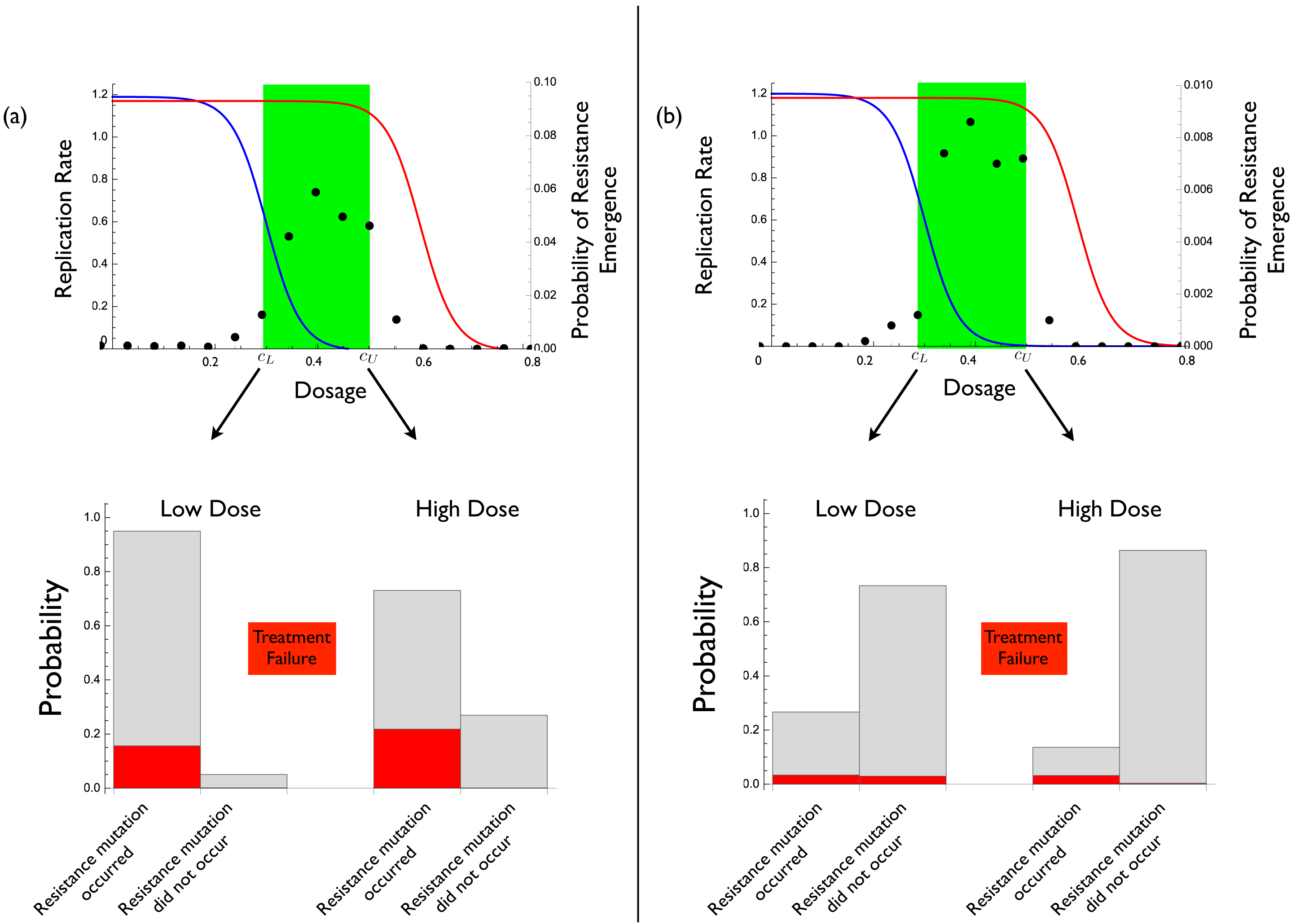
The effect of drug concentration on resistance emergence and treatment failure. (a) The dose-response curves for the wild type in blue (*r*(*c*) = 0.6(1 − tanh(15(*c* − 0.3)))) and the resistant strain in red (*r*_*m*_(*c*) = 0.59(1 − tanh(15(*c* − 0.6)))) as well as thetherpeutic window in green. Dots indicate the probability of resistance emergence. Probability of resistance emergence is defined as the fraction of 5000 simulations for which resistance reached a density of at least 100 (and thus caused disease). Parameter values are *P*(0) = 10, *I*(0) = 2, *α* = 0.05, *δ* = 0.05, *κ* = 0.075, *μ* = 10^−2^, and *γ* = 0.01. Bar graphs: the probability that a resistant strain appears by mutation is indicated by the left-hand grey bars for each drug concentration (the right-hand grey bar is the probability that a resistant strain does not appear). The probability of treatment failure for a specific drug dose is the sum of the red bars for that dose. (b) Same as panel (a) but with mutation rate decreased to *μ* = 10^−3^.

